# LINE-1 ribonucleoprotein complexes bind DNA to enable nuclear entry during mitosis

**DOI:** 10.1101/2024.10.01.616040

**Authors:** Sarah Zernia, Farida Ettefa, Srinjoy Sil, Cas Koeman, Joëlle Deplazes-Lauber, Marvin Freitag, Liam J. Holt, Johannes Stigler

## Abstract

Long Interspersed Nuclear Element-1 (LINE-1) is an autonomous retrotransposon that makes up a significant portion of the human genome, contributing to genetic diversity and genome evolution. LINE-1 encodes two proteins, ORF1p and ORF2p, both essential for successful retrotransposition. ORF2p has endonuclease and reverse transcription activity, while ORF1p binds RNA. Many copies of ORF1p assemble onto the LINE-1 RNA to form a ribonucleoprotein complex (RNP). However, the functional role of RNPs in the LINE-1 life cycle is unclear. Using reconstitution assays on DNA curtains, we show that L1 RNPs gain DNA binding activity only when RNA is super-saturated with ORF1p. In cells, L1 RNPs bind to chromosomes during mitosis. Mutational analysis reveals that DNA binding is crucial for nuclear entry and LINE-1 retrotransposition activity. Thus, a key function of ORF1p is to form an RNP that gains access to the genome through DNA binding upon nuclear envelope breakdown.

## Introduction

Transposable elements are genetic parasites that can move within the genome^1^. The long interspersed nuclear element (LINE-1, L1) is the only autonomously active retrotransposon^2^ in humans and accounts for about 17% of our genome^3,4^. The amplification of L1 in the human genome is one of the main drivers for genetic diversity and human evolution^5–7^, but insertion into essential genes can also have severe effects on the host organism, contributing to the development of diseases such as neurodegeneration and cancer^8,9^. Thus, it is of great interest to understand the mechanisms underlying retrotransposition.

LINE-1 encodes two proteins, ORF1p and ORF2p^10^, that drive a copy-and-paste retrotransposition process (**Figure 1A**). While both proteins are required for retrotransposition of LINE-1, the role of ORF1p in the LINE-1 lifecycle is poorly understood^11,12^. ORF2p is an endonuclease and reverse transcriptase that catalyzes the insertion of a new copy of L1 into the host genome^2,13^. ORF1p has a low-nanomolar affinity for RNA and has been proposed to be an RNA chaperone^14–16^. During translation, multiple copies of ORF1p bind the L1-mRNA and form a ribonucleoprotein complex (RNP) which has been suggested to protect L1 RNA from degradation^12,17,18^. This RNP formation is essential for transposon activity in vivo^19,20^, but the functional role of it remains unclear.

**Figure 1.**
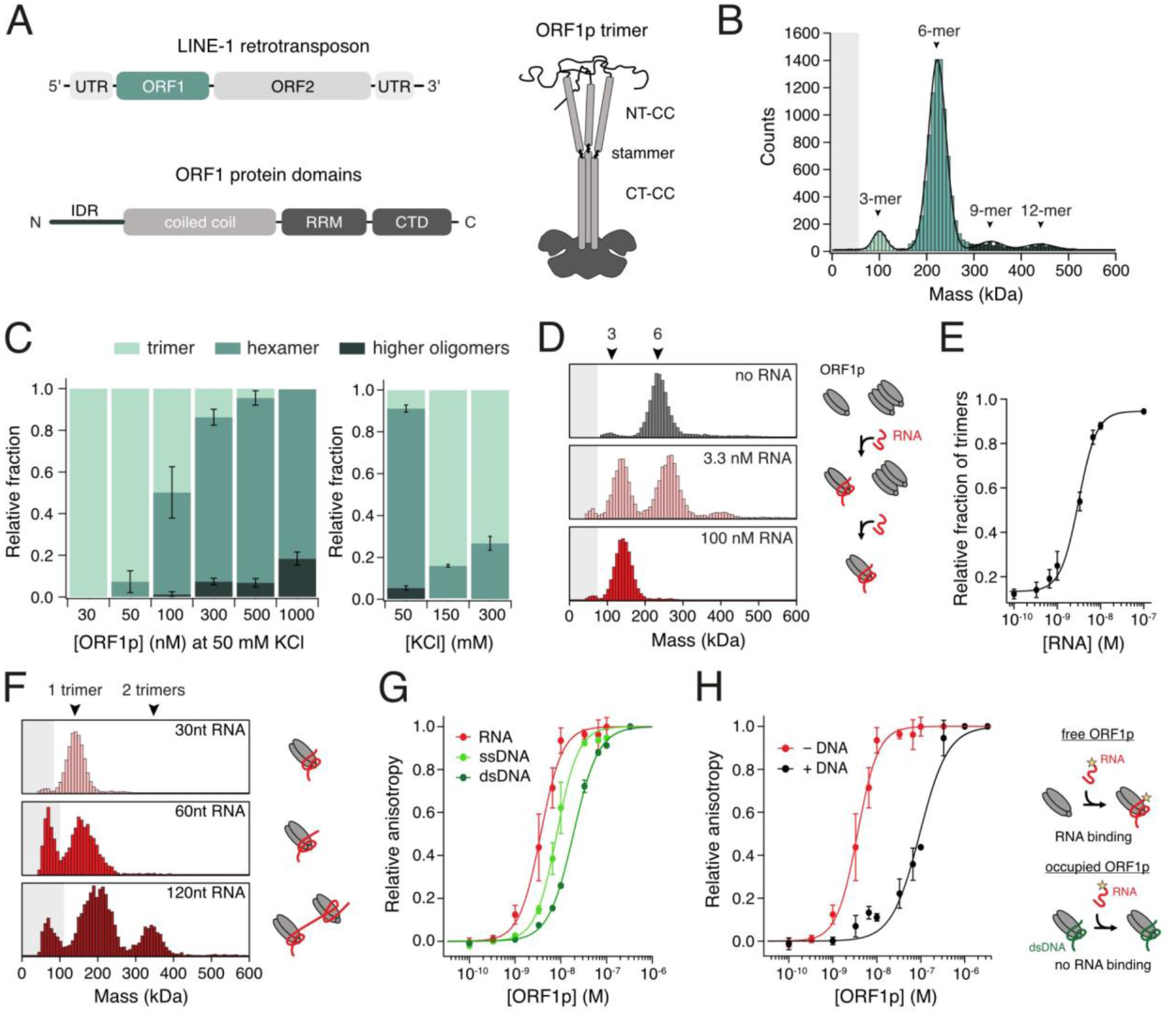
ORF1p forms oligomers that are modulated by RNA. **(A)** Top: Schematic representation of LINE-1 retrotransposon coding for ORF1p and ORF2p. Bottom: Domain composition of ORF1 protein containing an N-terminal intrinsically disordered domain (IDR), a long coiled coil followed by an RNA-recognition motif (RRM) and a C-terminal domain (CTD). Right: Schematic representation of the ORF1p homotrimer. The stammer (pos. 91-93) divides the coiled coil in N-terminal (NT-CC) and C-terminal coiled coil (CT-CC). **(B)** Mass photometry of 300 nM ORF1p at 50 mM KCl. The hexamer is the dominant species. Grey shading illustrates the detect limit. **(C)** Left: Mass photometry of ORF1p concentration series at 50 mM KCl. With higher protein concentrations, more and larger oligomers are formed. Right: Mass photometry of KCl series at 300 nM ORF1p. Oligomerization is reduced at higher salt concentrations, n=3, error bars represent SEM. **(D)** Mass photometry of 300 nM ORF1p mixed with increasing concentrations of 30 nt RNA. RNA presence leads to a dissociation of the hexamer. **(E)** Relative fraction of trimers at increasing RNA concentrations fitted with a Hill equation, h = 2.1 ± 0.3, n=5, error bars represent SEM. **(F)** Mass photometry of 300 nM ORF1p mixed with 100 nM of RNA of different lengths. 120 nt but not 60 nt RNA allows two ORF1p trimers to bind simultaneously. **(G)** Fluorescence anisotropy of ORF1p binding to 6-FAM-labeled 30 nt RNA, 30 nt ssDNA or 30 bp dsDNA. ORF1p has higher affinity for RNA than ssDNA or dsDNA. The binding curve was fitted with a Hill equation, n=3, error bars represent SEM. **(H)** Binding curves comparing binding of ORF1p to RNA (same as in (G)) versus ORF1p that was first saturated with 100 nM dark dsDNA and then bound to 1 nM 6-FAM-labeled 30 nt RNA (black). ORF1p cannot bind multiple nucleic acid molecules in parallel.

ORF1p is composed of an N-terminal intrinsically disordered domain (IDR), followed by a long coiled-coil (CC) domain and a globular C-terminal domain (CTD) containing the RNA recognition motif (RRM, **Figure 1A**)^21–24^. IDRs often facilitate multivalent interactions by electrostatic or also hydrophobic interactions^25–27^ and ORF1p IDR was shown to be required for condensate formation^19,20^. Additionally, the charge and position of a basic patch within the IDR are crucial for condensate formation^19^ and retrotransposition activity in cells^12,19,28,29^.

The ORF1p CC consists of seven heptad repeats that are interrupted by a three-amino acid “stammer motif” at position 91-93 that has been proposed to increase CC flexibility^28,30^. Interaction of the CC domains facilitates formation of a dumbbell-shaped ORF1p homotrimer^11,21,23^, which has previously been resolved by crystallography and NMR spectroscopy in parts^28,31,32^. However, ORF1p is also known to build higher oligomeric states^33,34^ of unclear structure; determining the nature of trimer-trimer interactions is crucial to resolve this question. We have previously studied the processes of phase separation and condensation of ORF1p in vitro and in human cells, respectively, and established condensate formation as critical for L1 retrotransposition^19^. However, the role of ORF1p oligomerization in L1 RNP assembly and retrotransposition remains unclear.

Oligomerization has been shown to affect ORF1p’s interactions with nucleic acids^33^, which is of great interest given the requirement of L1 proteins to interact with both RNA and DNA during retrotransposition. Beyond its high affinity for RNA^33,35^, ORF1p has also been shown to bind to DNA in both bulk^33^ and single-molecule experiments^34,36^. This raises the possibility that ORF1p could be involved in DNA target site recognition or priming. Previously, DNA-binding activity was investigated with ORF1p alone, but never within assembled RNP complexes. As a result, it is unknown whether this DNA-binding capability is restricted to the isolated protein or could be extended to the ORF1p-RNP and whether ORF1p has another role in retrotransposition besides RNA protection.

Here, we combine single-molecule DNA curtain^37^ technology, mass photometry^38^ and fluorescence anisotropy with high-resolution cell imaging to reveal the interplay between protein oligomerization and nucleic acid-binding, RNP assembly and its localization to DNA, and L1 retrotransposition. We show that ORF1p forms multimers of trimers, which is driven by electrostatic interactions and tuned by RNA binding. Furthermore, we reconstituted RNPs by mixing L1 RNA and ORF1p, and determined that these RNPs bind to DNA. By altering RNP composition, we show that excess ORF1p is required to generate dsDNA-interacting RNPs, consistent with the high stoichiometry of ORF1p observed in cells^12^. Mutational studies suggest a connection between oligomerization behavior and stable DNA binding of RNPs. We present that L1 RNPs show co-localization and correlated positioning with mitotic chromatin in cells and only can gain access to the nucleus to successfully complete retrotransposition if they interact with DNA. Together, our study establishes direct DNA binding as a new function for L1 RNPs. Thus, we provide insights into how ORF1p oligomerization and nucleic acid interactions are interdependent in RNP formation as well as target site recognition and determine a crucial role for ORF1p-RNPs to initiate LINE-1 retrotransposition.

## Results

### RNA modulates ORF1p oligomerization

We investigated the oligomerization states of purified human ORF1p using single-molecule mass photometry^38–40^. First, we analyzed the mass distribution of 300 nM of ORF1p at 50 mM KCl and detected only minor amounts of trimer but a high proportion of hexamers and significant peaks for higher oligomeric forms (**Figure 1B, S1A**). Upon testing oligomerization ability under different conditions, we discovered that with lower protein concentration, the proportion of hexamers decreased while the proportion of trimers increased (**Figure 1C**). The same trend was observed with increasing salt concentrations. This suggests that oligomerization is driven by both protein concentration and electrostatic interactions. ORF1p monomers were hardly detected which may be due to the detection limit of the mass photometry device (about 40 kDa), but could also suggest that trimerization is very stable and preferred^32,33^.

Next, we sought to understand the impact of RNA on this oligomerization behavior. We used a 30 nt unstructured RNA derived from the 5’-UTR of LINE-1. This RNA length was shown before to be sufficient for one trimer to bind^33^. With increasing RNA concentrations, ORF1p oligomers dissociated into trimeric species (**Figure 1D, S1A)**, suggesting that trimer-trimer interactions are disturbed by RNA binding. Upon closer inspection of the RNA-induced dissociation of ORF1p hexamers, we found that this process can be modeled by a Hill equation with a Hill coefficient of 2.1 ± 0.3, compatible with the cooperative generation of two trimers (**Figure 1E**). RNA binding also induced a mass shift of both the trimer and the hexamer, suggesting that both species are able to bind to RNA.

In agreement with previous studies^33^, our results demonstrate that RNA concentration can tune ORF1p oligomerization. We speculated that this tunability might be necessary for forming stable RNPs. Under equimolar conditions (100 nM ORF1p trimer + 100 nM RNA), only the trimer peak was observed in mass photometry, suggesting that the 30 nt RNA provided space for only one ORF1p trimer to bind. We then increased the RNA length to 60 nt and still observed only one trimer on the RNA. However, at 120 nt length, we observed besides the peak for one trimer also a peak for two ORF1p trimers, which we believe to associate with the same RNA molecule (**Figure 1F**). Thus, while 30 nt RNA are sufficient for one ORF1p trimer to bind^32,33^, an RNA length between 60 and 120 nt is required to accommodate two trimers.

To support these findings, we modeled RNA binding of an ORF1p trimer using AlphaFold 3^41^ (**Figure S1B**). The RNA binding site is located at a strongly positively charged cleft between the RRM and the CTD, which can open and close upon substrate binding^24,31,32^. In five computed models of the trimer, the RNA was wrapped around the three CTDs and interacted with each binding cleft of the individual monomers as postulated previously^32,42^. Remarkably, we found that the 30 nt RNA is fully occupied by one trimer. Thus, it would require a short linker between two binding sites to avoid steric clashes of two trimers. This could be a possible explanation for our observation that two ORF1p trimers require more than 60 nt to bind.

### RNA and DNA compete for the same binding site on the ORF1p trimer

The exact binding mode of RNA to the ORF1p trimer is unknown as no nucleic acid-bound structure is available. The curved and narrow binding cleft suggests a preference for flexible and unstructured nucleic acids^32^. We determined the binding affinities of ORF1p to different nucleic acid substrates using fluorescence anisotropy. To this end, we titrated unlabeled ORF1p to fluorescently labeled 30 nt RNA, 30 nt ssDNA or 30 bp dsDNA. The resulting binding curves (**Figure 1G**) showed the highest affinity to RNA (K_D_ = 3.9 ± 1.1 nM), followed by ssDNA (K_D_ = 8.3 ± 1.8 nM) and dsDNA (K_D_ = 18.9 ± 1.3 nM), in agreement with previous studies^32,33^. Additionally, we tested binding of ORF1p to alternative nucleic acid substrates and found that there is no significant difference between RNA and an RNA/DNA hybrid (K_D_ = 4.7 ± 0.6 nM) or a poly(A) (K_D_ = 6.0 ± 1.2 nM, **Figure S1D**). Note that RNA affinity was slightly reduced at higher salt concentrations (**Figure S1E**).

As we saw reduced binding affinity of ORF1p towards dsDNA, we wondered if this implies that DNA binds at a different site or with a different binding mode than RNA. To determine whether ORF1p oligomerization is altered upon binding of dsDNA, we repeated the mass photometry experiments in presence of increasing concentrations of 30 bp dsDNA. Interestingly, we observed that dsDNA induces hexamer dissociation in a manner comparable to RNA (**Figure S1C**) suggesting a similar binding mode for RNA and DNA. Following up on this observation, we wondered if RNA and dsDNA bind to the same site on ORF1p. To this end, we saturated ORF1p with 100 nM dsDNA and tested subsequent binding to 1 nM labeled RNA (**Figure 1H**). The shift of the binding curve demonstrates a drop in RNA affinity by 100-fold, indicating that ORF1p that is occupied by dsDNA is unable to bind to RNA. Similar results were found when first saturating with 100 nM RNA and then adding 1 nM labeled dsDNA (data not shown), implying that both nucleic acids have comparable binding modes and compete for the same binding site on ORF1p.

### Excess protein enables ORF1p-RNP binding to DNA

So far, we have investigated ORF1p binding to small oligos to study the details of ORF1p trimer interactions with nucleic acids. However, in a naturally occurring ORF1p-RNP, many ORF1p trimers co-translationally assemble onto a long LINE-1 mRNA. Therefore, we used ORF1p bound to a 2 kb fragment of the L1 RNA as a model for our subsequent studies. Interactions between ORF1p and these longer RNAs led to the formation of RNP complexes, which we found in a previous study to be essential for LINE-1 retrotransposition^19^. We hence hypothesized that RNPs might bind to dsDNA.

To test this idea, we employed single-molecule DNA curtains, which consist of arrays of lambda phage DNA (λ-DNA) stretched between two chromium barriers in a microfluidic flow cell (**Figure 2A**)^37,43^. In this system, each individual dsDNA molecule can be directly visualized and binding of fluorescently labeled molecules can be monitored in real time using TIRF microscopy.

**Figure 2.**
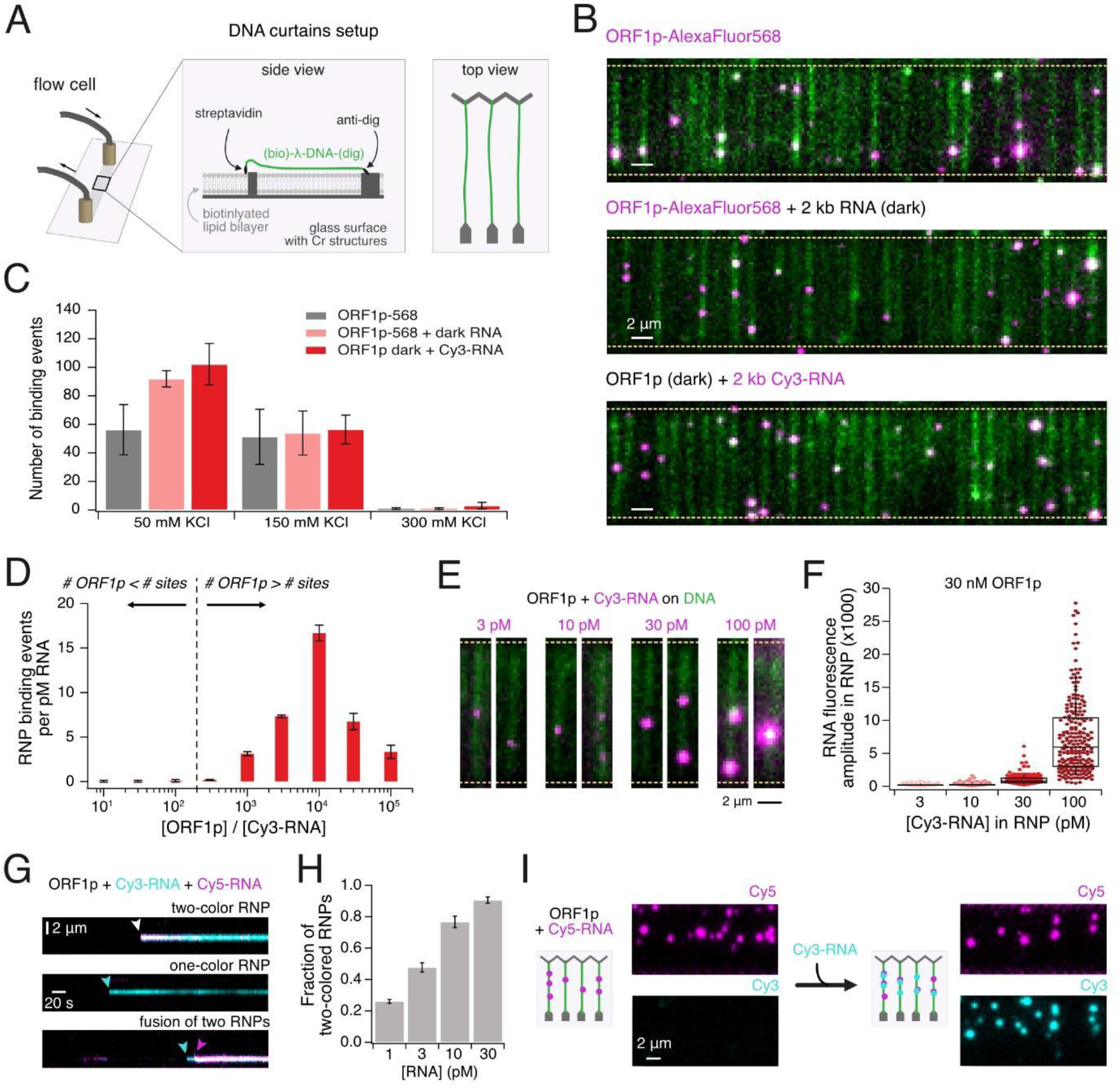
RNPs formed by ORF1p and a 2 kb L1 RNA fragment bind efficiently to double stranded DNA curtains. **(A)** Schematic representation of the DNA curtains experimental setup. Lipid-bound λ-phage DNA is introduced into a custom-built flow cell that contains chromium structures as diffusion barriers (Cr barrier). Flow causes the λ-phage DNA to stretch across the chromium barrier and anchor to a second chromium structure, thus holding the DNA in a stretched configuration. Other molecules can then be introduced to the flow cell to assess binding to DNA. **(B)** TIRF microscopy of ORF1p or ORF1p-RNPs (magenta) binding to λ-DNA (green). Yellow dashed lines illustrate Cr barrier position. Top: 30 nM Alexa Fluor 568-labeled ORF1p; middle: 30 nM Alexa Fluor 568-labeled ORF1p pre-incubated with 30 pM 2 kb RNA (i.e. 1000:1 ORF1p:RNA); bottom: 30 nM dark ORF1p pre-incubated with 30 pM 2 kb Cy3-labeled RNA. **(C)** Average number of binding events on DNA curtains at different KCl concentrations, n≥3, error bars represent the SEM. **(D)** Weighted average of RNP binding events per RNA concentration at 150 mM KCl. The dashed line marks 200x excess of ORF1p, which is predicted to saturate all ORF1p trimer binding sites on RNA assuming a minimum spacing of 30 nucleotides between trimers. If the L1 RNA is saturated with ORF1p trimers, no binding occurs. Once the L1 RNA is saturated for direct trimer binding, the RNP starts to bind DNA. n=3, error bars represent the SEM. **(E)** TIRF microscopy images of ORF1p-RNPs (magenta) containing different amounts of Cy3-RNA binding to DNA curtains (green). The brightness of RNPs increases with increasing RNA concentration. **(F)** Fluorescence amplitudes of ORF1p-RNPs binding to DNA curtains at 150 mM KCl. 30 nM dark ORF1p were mixed with increasing amounts of Cy3-RNA leading to an increased fluorescence intensity, n≥3. **(G)** Kymograms of DNA-binding events of 30 nM ORF1p pre-mixed with 15 pM Cy3-RNA (cyan) and 15 pM Cy5-RNA (magenta): RNP containing two colors of RNA (top), RNP containing a single RNA color (middle), fusing RNPs of two different colors (bottom). **(H)** Fraction of RNPs containing two RNA colors at different total RNA concentrations. Cy3-RNA and Cy5-RNA were added at an equimolar concentration. n=4, error bars represent the SEM. **(I)** DNA-bound ORF1p-RNPs can recruit additional RNAs. ORF1p was pre-incubated with Cy5-RNA (magenta), RNPs were confirmed to bind to DNA, and no fluorescence was observed in Cy3-channel. Cy3-RNA (cyan) was then added to the flow cell and was recruited to already bound RNPs. Cy3-fluorescence overlaps with Cy5-fluorescence.

We wondered how ORF1p would interact with DNA either alone or as an RNP complex. First, we added fluorescently labeled ORF1p to the DNA curtains in the absence of RNA and observed robust protein binding to the DNA (**Figure 2B**, top; **Figure S2A**). Next, we reconstituted RNPs by pre-incubating fluorescently labeled ORF1p with unlabeled 2 kb RNA for 5 minutes, which we then applied to the DNA curtains. These RNPs also bound robustly to the DNA (**Figure 2B**, middle). The same observation was made when labeling the RNA instead of the protein within the RNP (**Figure 2B**, bottom). This result indicates that although one ORF1p trimer cannot bind to RNA and DNA simultaneously (**Figure 1H**), ORF1p-RNPs can. Additionally, this binding is very stable as more than 60 % of the bound RNPs remained bound for more than 5 min (**Figure S2C**). We next quantified binding of ORF1p alone and ORF1p-RNPs at different salt concentrations. At 50 mM KCl, the RNP exhibited enhanced binding compared to ORF1p alone, while at 150 mM KCl, the RNP and protein alone displayed similar binding counts (**Figure 2C**). No binding was observed for either species at 300 mM KCl. Thus, we conclude that RNP binding to DNA relies upon electrostatic interactions.

To determine the likelihood of ORF1p forming an RNP in the presence of RNA, we labeled both protein and RNA, and quantified the frequency of co-localization between the two signals. Strikingly, in almost all cases, ORF1p was part of an RNP (**Figure S2D**) indicating that ORF1p preferred to bind DNA in the form of RNPs when in the presence of RNA, even though ORF1p was in high stoichiometric excess. Cy3-RNA did not bind on its own to DNA curtains (**Figure S2B**, top). Therefore, we used RNPs containing labeled RNA and dark ORF1p for the following experiments to ensure the observed binding events were only attributed to RNPs.

To evaluate whether ORF1p-RNPs have a preferred binding site on DNA, we analyzed binding positions and found that they correlate, to some extent, with the known LINE-1 ORF2p target site TTTTTAA^13,44–47^ and in general with A/T-rich DNA regions (**Figure S2F**). This suggests that ORF1p-RNPs could play a role in target site recognition and LINE-1 stabilization on genomic DNA.

We next investigated the optimal ratio of protein to RNA for DNA binding. We determined the number of binding events while keeping either RNA or ORF1p concentration constant and varying the other RNP component. For better data interpretation, we normalized the data with the respective RNA concentration as the RNA carries the label (**Figure 2D**, unnormalized data **Figure S2E**). Assuming that the footprint of one ORF1p trimer on RNA is 30 nt, the 2 kb RNA provides a maximum of 68 binding sites, which means that all binding sites would be occupied by a 200-fold molar excess of protein. Interestingly, we did not observe any RNP binding to DNA with an ORF1p:RNA stoichiometric excess below 200-fold (**Figure 2D**), a ratio where each ORF1p nucleic acid-binding cleft is occupied by RNA and can no longer bind to DNA, similar to the scenario in **Figure 1H**. In support of this, when we mixed ORF1p in 1:1 molar ratio with short RNA oligos, we also did not observe any DNA binding (**Figure S2B**, top). In contrast, when the ORF1p concentration was in 1000-fold excess, RNP binding to DNA was robust and further increased with higher protein concentrations (**Figure 2D**). These results suggest that, to bind DNA, RNPs must contain an excess of ORF1p trimers, beyond the amount required to fully saturate the L1 RNA molecule.

### ORF1p-RNPs are variable in composition

Upon testing DNA binding of RNPs formed with various concentrations of labeled RNA, it became apparent that RNPs become brighter with increasing RNA concentrations (**Figure 2E–G**). This indicates that ORF1p-RNPs can, in principle, incorporate more than one RNA molecule. To assess this hypothesis, we mixed ORF1p with two differently colored RNAs. Not only did this yield two-colored RNPs (**Figure 2G**), but the fraction of two-colored RNPs also increased to 90.5 ± 0.1% as we increased the RNA concentration to 30 pM at an equal ratio of the two colors (**Figure 2H**). Furthermore, RNPs that are already bound to DNA can recruit additional RNA molecules (**Figure 2I**), suggesting that DNA-binding competent RNPs are still multivalent. Supportive to that, we also observed fusion of RNPs. We increased the incubation time of protein and RNA from 1 minute to 10 or 30 minutes and found a strong increase in fluorescence after a 30 minutes incubation (**Figure S2G**) demonstrating that with more time, larger RNPs can form. The same was observed when mixing differently colored pre-formed RNPs before loading them onto the DNA. In 56.5 ± 11.3 % of the cases we found two-colored fused RNPs (**Figure S2H**).

### The ORF1p N-terminus is required for the DNA-binding activity of RNPs

As RNP formation is influenced by both ORF1p oligomerization and nucleic acid interaction, we wondered whether we could assign these activities to distinct regions within the ORF1p structure. To this end, we created ORF1p variants with a mutation of the N-terminal basic patch (K3A/K4A-ORF1p), a deletion of the CC stammer motif (ΔStammer-ORF1p) or an elongated N-terminus (His-TEV-ORF1p).

We first started by determining the RNA binding affinity of the three variants and found only a slight increase in K_D_ compared to wildtype (**Figure 3B**). Next, we investigated their oligomerization using mass photometry and found no change in oligomerization of K3A/K4A-ORF1p, but, surprisingly, a stabilization of the hexamer and higher-order oligomers for the ΔStammer-ORF1p (**Figure 3C**). This stabilization is even more pronounced at 150 mM KCl (**Figure 3D**), which suggests that the deletion of the stammer, and therefore a change in the flexibility of the CC^28^, stabilizes inter-trimer interactions. This is in line with previous studies identifying the CC domain as the main trimer-trimer interaction point^34,36^. The N-terminally elongated variant (His-TEV-ORF1p) displayed strongly impaired oligomerization (**Figure 3C+D**). It appeared that even trimers could not form correctly, as monomers were observed in 30 % of all counts. This is likely a lower-bound estimate of the fraction of monomers, as the mass of an ORF1p monomer (40 kDa) is close to the lower detection limit of the mass photometer^48^. With an N-terminal elongation, contact points between N-terminal amino acids and the CC domain that are essential for trimer-trimer interactions^20^ are probably lost, leading to impaired oligomerization.

**Figure 3.**
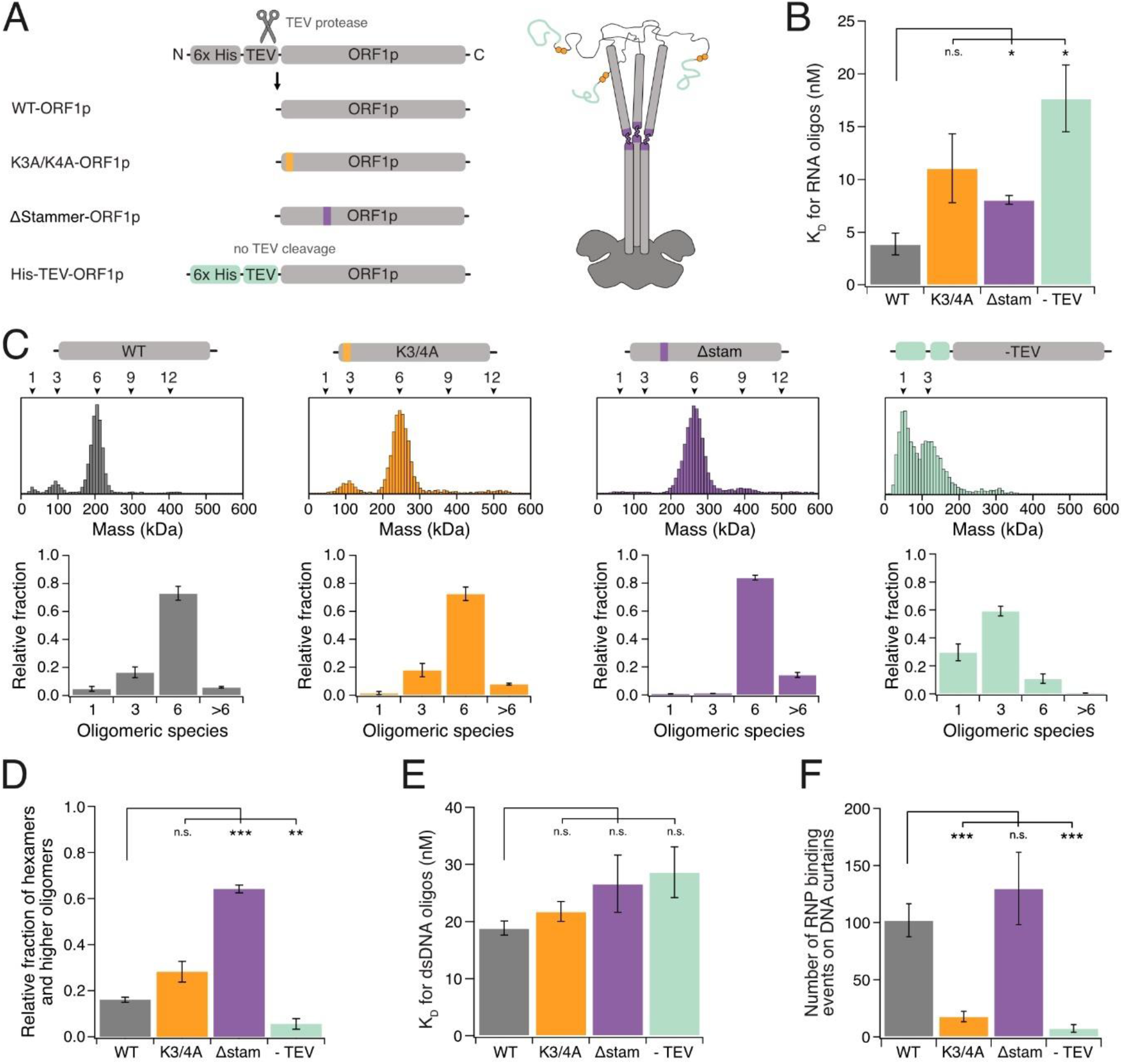
ORF1p variants exhibit different oligomerization and nucleic acid-binding properties. **(A)** Schematic of ORF1p variants showing the location of the mutations within the homo-trimer. WT-ORF1p, K3A/K4A-ORF1p and ΔStammer-ORF1p were cut by TEV protease to release a free N-terminus. His-TEV-ORF1p was not cut by TEV protease resulting in an elongated N-terminus. **(B)** Dissociation constants of RNA binding of ORF1p variants determined by fluorescent anisotropy. n=3, error bars represent the SEM, significance determined by Student’s t-test. **(C)** Mass photometry of ORF1p variants. Top: Histogram of molecular masses. Bottom: Relative fraction of oligomeric species (1: monomer, 3: trimer, 6: hexamer, >6: higher oligomeric forms). ΔStammer-ORF1p shows better oligomerization, His-TEV-ORF1p oligomerization is strongly impaired. **(D)** Relative fraction of hexamers and higher-order oligomers of ORF1p variants (300 nM) at 150 mM KCl determined by mass photometry. ΔStammer-ORF1p shows significantly more oligomers, His-TEV-ORF1p shows significantly less oligomers than WT. n=3, error bars represent the SEM. Significance determined by Student’s t-test. **(E)** Dissociation constants for ORF1p variants binding to short dsDNA oligos was determined by fluorescence anisotropy. ORF1p variants have similar affinity to dsDNA as WT. n=3, error bars represent the SEM. Significance determined by Student’s t-test. **(F)** Average number of binding events of 30 nM ORF1p variants and 30 pM Cy3-RNA on DNA curtains at 50 mM KCl. K3A/K4A-ORF1p and His-TEV-ORF1p bind significantly less. n≥4, error bars represent the SEM. Significance determined by Student’s t-test.

Next, we wondered how RNPs formed from these ORF1p variants with 2 kb RNA interact with DNA curtains. All protein variants had a comparable DNA-binding affinity in fluorescence anisotropy (**Figure 3E**). However, RNPs formed by His-TEV-ORF1p were hardly able to bind to DNA curtains (**Figure 3F**). This result implies that the N-terminal elongation does not impair DNA binding of the protein per se but particularly the DNA binding of the RNP, most likely because of an altered RNP composition due to a different oligomerization behavior of the mutant. ΔStammer-ORF1p-RNPs showed similar binding compared to WT, but K3A/K4A-ORF1p-RNPs bound significantly worse than WT-ORF1p-RNPs on DNA curtains (**Figure 3F**) even though hexamer formation and RNA binding was not affected by this mutation. Basic patches, especially in IDRs, form electrostatic interactions with the phosphate backbone of RNA^49^ and long RNA molecules can themselves modulate the chemical environment of this disordered regions to favor ribonucleoprotein complex assembly^50–52^. As K3A/K4A-RNPs are less DNA-binding competent, we speculate that the basic patches of multiple IDRs form a second binding site for nucleic acids, which then promote further RNP assembly. This would also explain the reduced DNA-binding of His-TEV-ORF1p-RNPs as in this variant the position of the basic patch is shifted^12,19,28,29^, possibly preventing the formation of this additional nucleic acid binding site. Together, these results show that the N-terminus of ORF1p is critical for the DNA binding of ORF1p-RNPs.

### The DNA-binding activity of ORF1p is necessary for LINE-1 nuclear entry and retrotransposition

Given the ability of ORF1p to bind DNA both on its own and within the context of an RNP, we wondered if we could observe binding of ORF1p to DNA in cells and if this activity is relevant to LINE-1 retrotransposition in cells. We used a previously described Tet-inducible, active, endogenous-like L1 expression construct in which ORF1p was fused at its C-terminus to a HaloTag (ORF1p-Halo)^53^ (**Figure 4A**). This enabled visualization of ORF1p in live cells by labeling with fluorescent Janelia Fluor HaloTag Ligands JFX549^54^. The construct also contained a reporter to assess the ability of this engineered construct and its variants to complete the full L1 retrotransposition cycle. The reporter consisted of a GFP cassette interrupted by an antisense γ-globin intron (GFP-AI) in the 3’ UTR of our construct^11,55,56^. The antisense disrupts the GFP open reading frame such that transcription, splicing, and successful retrotransposition of this spliced L1 mRNA is required for subsequent expression of an uninterrupted GFP coding sequence from a novel insertion site. This reporter enabled us to relate changes in ORF1p behavior upon mutation to the completion of the L1 retrotransposition life-cycle.

**Figure 4.**
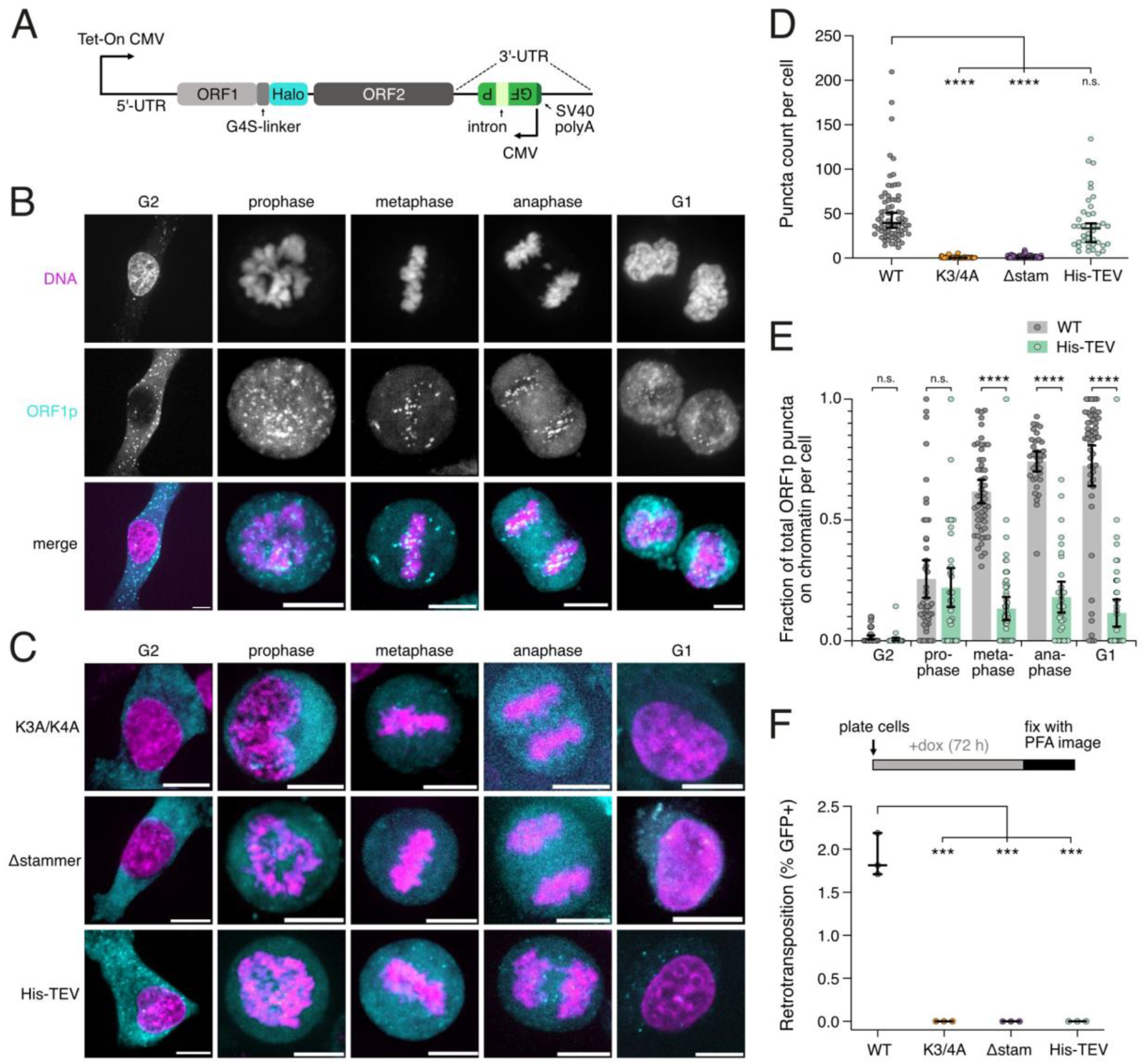
Binding of ORF1p puncta to mitotic chromatin is required for LINE-1 nuclear entry and retrotransposition. **(A)** Schematic of Tet-inducible LINE-1 expression construct with a GFP-antisense intron (GFP-AI) retrotransposition reporter. ORF1p is tagged with a Halo-tag that can be visualized using a Halo ligand conjugated with JFX549 or JFX554. GFP is expressed in the opposite orientation and interrupted by a gamma-globin intron such that transcription, removal of the intron by splicing, and subsequent retrotransposition is required for GFP expression. **(B)** WT-ORF1p puncta bind mitotic chromatin following nuclear envelope breakdown and associate with chromosomes throughout mitosis. HeLa cells expressing WT-ORF1p-Halo were treated with doxycycline for 6-12 hours, then treated with SiR-DNA to stain DNA and JFX549 to label the Halo-tag, then imaged. The DNA signal (top), WT-ORF1p-Halo signal (middle), and merged image (bottom) are shown for various cell cycle stages (left to right). Representative maximum intensity projections of Z-stacks are shown. **(C)** K3A/K4A-ORF1p and ΔStammer-ORF1p were unable to form puncta while His-TEV-ORF1p formed puncta that did not bind mitotic chromatin. Representative images of each mutant (top to bottom) for various cell cycle stages (left to right) are shown. The same imaging paradigm was performed as in **(B). (D)** WT- and His-TEV-ORF1p robustly form puncta, but the K3A/K4A-ORF1p and ΔStammer-ORF1p did not. Significance determined by Student’s t-test. N>30 per cell cycle stage, n=3. **(E)** Quantification of the colocalization between ORF1p puncta and chromatin at various stages of the cell cycle for WT-ORF1p and His-TEV-ORF1p. WT-ORF1p puncta show increased association with mitotic chromatin with increased compaction during mitosis relative to His-TEV-ORF1p. The number of puncta colocalized with chromatin was divided by the total number of puncta in each cell. Significance determined by multiple Student’s t-tests. N>30 per cell cycle stage, n=3. **(F)** DNA-binding activity of ORF1p is necessary for LINE-1 retrotransposition. Top: Cells were continuously treated with doxycycline for 72 hours and retrotransposition rates were calculated using the GFP-AI reporter. Bottom: The percentage of cells expressing GFP (GFP+) obtained by imaging were scored for each ORF1p mutant in addition to WT. Significance determined by Student’s t-test. N>400, n=3.

Consistent with our prior results, formation of ORF1p puncta was observed after around 6 hours of induction of our expression system with doxycycline. We compared WT-ORF1p puncta formation to that of the mutants we had already assessed in vitro: K3A/K4A-, ΔStammer- and His-TEV-ORF1p (**Figure 4B-D**). At early time points, WT- and His-TEV-ORF1p formed a similar number of puncta while K3A/K4A-ORF1p and ΔStammer-ORF1p did not form puncta at all (**Figure 4D**). The WT-ORF1p puncta were mostly localized to the cytoplasm but sometimes also to the nucleus (**Figure 4B)**. Interestingly, while there were minimal nuclear puncta in G2 prior to nuclear envelope breakdown, we observed substantial co-localization of ORF1p with chromatin upon nuclear envelope breakdown in prometaphase. This colocalization increased during metaphase and anaphase, and persisted into the subsequent G1 (**Figure 4B, S3A+D**). This is consistent with prior reports of ORF1p nuclear entry during mitosis as a result of nuclear envelope breakdown, and retention in the daughter nuclei due to nuclear envelope reassembly post-division^11^. We additionally noted correlation between the positions of the ORF1p puncta and the mitotic chromatin, suggesting binding to the DNA as a mechanism for LINE-1 nuclear entry.

We next analyzed the behavior of our N-terminal elongated His-TEV-ORF1p mutant. Expression of His-TEV-ORF1p produced puncta that had greatly reduced colocalization with mitotic chromatin, and greatly diminished nuclear puncta in G1 relative to WT (**Figure 4C+E, S3B+C+F**). To assess the role of ORF1p’s DNA-binding activity in LINE-1 retrotransposition, we induced WT and mutant ORF1p expressing HeLa cells with doxycycline for 72 hours, and looked for GFP expression indicating successful retrotransposition events (**Figure 4A**). While WT-ORF1p cells underwent retrotransposition at a rate of about 2 % of cells, which is on par with our prior observations^19^, no retrotransposition events were observed among the cells expressing His-TEV-ORF1p or the other two mutants (**Figure 4F**). These data support the hypothesis that the DNA-binding activity of ORF1p is necessary for LINE-1 retrotransposition. Taken together, the results from our cell experiments suggest a model where ORF1p binds DNA following nuclear envelope breakdown during mitosis to enable the LINE-1 RNP to access the genome and carry out retrotransposition.

## Discussion

### DNA binding of ORF1p-RNPs enables nuclear entry

Here, we described DNA targeting of LINE-1 RNPs comprised of RNA and ORF1p. Our reconstitution and single-molecule imaging significantly advance our understanding beyond previous studies of isolated ORF1p^33,34,36^. Our data indicates that while isolated ORF1p has the ability to bind DNA, its comparatively higher RNA affinity causes it to preferentially assemble into RNPs with RNA (**Figure 2**). As ORF1p binds co-translationally to L1 mRNA in cells (cis preference)^22,57^, we propose that this initial RNP formation could serve multiple functions in the cytoplasm including melting of RNA secondary structure as shown for other high-affinity RNA binders^58,59^, and protection of the L1 mRNA from the innate immune system^60–62^. After translation, the RNP must gain access to the nucleus to allow retrotransposition on genomic target sites. As the RNP complex is likely several hundred nanometers in diameter, it is too large to enter the nucleus through the nuclear pore^63,64^. As also reported for several retroviruses^65^, our study is consistent with a model where L1 gains access to the genome by binding chromosomes when the nuclear envelope is disassembled during mitosis^11^. The ability of L1 RNPs to directly bind DNA in vitro **(Figure 2)**, and the observation that these particles also bind mitotic chromatin **(Figure 4)**, suggests that the L1 RNP acts as a transport vehicle to bring L1 RNA into the nucleus, and into proximity to known integration sequences on DNA.

### LINE-1 RNPs evade mechanisms that mediate exclusion of cytoplasmic contents from the nucleus

During nuclear envelope breakdown it is crucial that cytoplasmic and nuclear contents stay separated. Mitotic chromosomes contain proteins, such as Ki-67 and BAF, that coat the outside of the chromosome^66^ to prevent DNA-interaction of cytoplasmic macromolecular objects^67^. Preventing invasive molecules like retrotransposons from gaining access to the genome is an essential host-defense mechanism. Given our observation of L1 RNPs co-localizing with mitotic chromatin, we speculate that L1 RNPs can overcome this Ki-67- and BAF-mediated nuclear exclusion. Perhaps, L1 RNPs have biophysical properties that allow them to partition into the perichromosomal layer formed by Ki-67 during mitosis^66,68,69^ or that enable association to dense chromatin networks that are formed by BAF during nuclear envelope reformation^67,70^.

### ORF1p may guide RNPs to L1 target sites

Our reconstitution experiments suggest that ORF1p-RNPs could play a role in L1 target site acquisition. Recently, a study structurally resolved ORF2p binding to the DNA target site showing a very tightly regulated recognition of the target sequence TTTTTAA^47^. For ORF1p, no sequence-specific nucleic acid-binding has been reported^35^. We observed only a slight target site preference, but ORF1p-RNPs did get enriched on A/T rich regions of DNA (**Figure S2F**). Therefore, we hypothesize that ORF1p guides the RNP towards A/T-rich regions, within which ORF2p then has a higher probability to specifically recognize the L1 integration target sequence.

Besides DNA binding, ORF1p is known for its nucleic acid chaperone activity, inducing DNA melting, strand annealing and strand exchange^14,33,36,71,72^. These processes are more likely to occur at A/T-rich regions where base pair interactions are weaker^73^. In accordance, we observed most ORF1p and ORF1p-RNP binding events on DNA at A/T-rich regions, and these binding events were very static (**Figure S2A**) suggesting that ORF1p may locally melt the DNA in our assay. Fluorescence anisotropy data further showed that ORF1p has a very high affinity for RNA/DNA hybrids (**Figure S1D**), raising the possibility that ORF1p is involved in hybridization of L1 RNA with the target site and other strand annealing processes during the reverse transcription mechanism^45,71^.

### ORF1p stoichiometry tunes L1 RNP function

Strikingly, ORF1p-RNPs have no DNA-binding activity until ORF1p has super-saturated the RNA. This is likely caused by the high affinity of ORF1p for RNA and the fact that it can only bind one nucleic acid molecule at a time (**Figure 1H**). Thus, ORF1p initially coats the RNA and all nucleic acid binding sites are occupied. However, with higher ORF1p excess, protein-protein interactions enable recruitment of additional ORF1p into the RNP, which then still have free nucleic acid binding sites. These super-saturated RNPs can now efficiently bind to DNA (**Figure 5**). This means that both efficient RNA binding by ORF1p and a high degree of ORF1p oligomerization are prerequisites for DNA targeting of L1. We speculate that this is a mechanism to prevent off-target activity of ORF1p. In this model, its high affinity for RNA might cause ORF1p to be sequestered by the general cellular RNA pool, but these sub-stoichiometric amounts of ORF1p on incorrect RNA targets would not be able to target DNA and would hence be relatively inert. Only ORF1p co-translationally assembled at high stoichiometry on the L1 mRNA would gain DNA-binding activity. Recent evidence suggests that ORF1p-RNPs indeed assemble predominantly in cis on the mRNA from which they were translated^19^.

**Figure 5.**
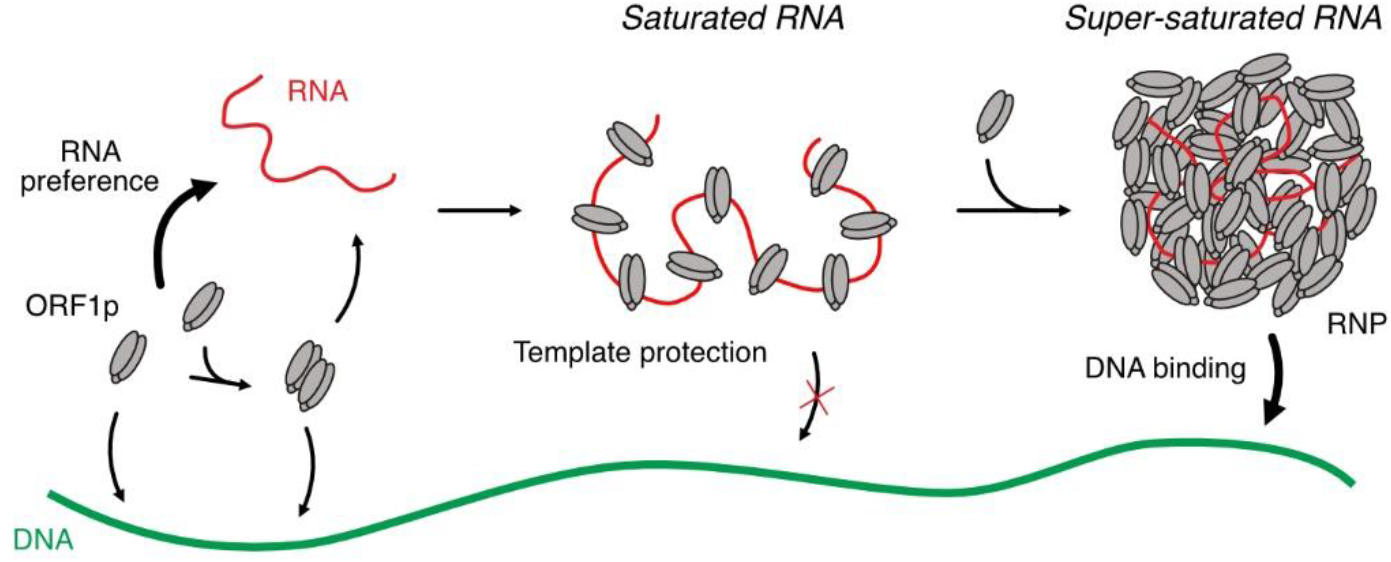
Model for ORF1p-RNP formation and DNA recognition. ORF1p forms homo-trimers and higher-order oligomers that can bind to DNA but have a clear preference for RNA. Upon RNA binding, ORF1p trimers occupy first all available binding sites on the RNA, which protects the RNA from degradation, but is unable to bind to DNA. Continued ORF1p accumulation leads to super-saturation, which is required to form a DNA-binding-competent RNP.

Together, our results suggest that ORF1p could confer distinct functions to the L1 RNP depending on stoichiometry. This provides a kinetic control of L1 RNP function during co-translational assembly. At the beginning of translation, excess RNA disrupts the formation of ORF1p trimer-trimer contacts and favors a configuration, where the LINE-1 RNA is sequestered in the RNA-binding cleft, perhaps providing protection against degradation, or avoiding detection by the innate immune system. With continued translation, ORF1p will super-saturate the L1 RNA, and excess nucleic acid binding sites render RNPs competent to bind to mitotic chromatin at potential integration sites. Thus, L1 condensates have the remarkable feature that a new function – DNA binding – emerges at a threshold stoichiometry of ORF1p. It will be interesting to see if other ribonucleoprotein condensates undergo functional changes when their component stoichiometries change.

## Acknowledgements

We thank Philipp Gerlof for initial experiments on His-TEV-ORF1p and Jeanny Probst for preparation of DNA templates for RNA synthesis. Thanks also to Daniel Bollschweiler, MPIB Martinsried as well as Gregor Witte and Karl-Peter Hopfner, Gene Center Munich for providing access to the mass photometer. This work was supported by the LMU-NYU Research Cooperation Program.

## Author contributions

S.Z performed DNA curtains, fluorescence anisotropy and mass photometry experiments. F.E. expressed and purified ORF1p-WT and ORF1p-K3A/K4A proteins, produced long RNA for DNA curtains measurements and performed cell experiments. S.S. cloned WT, K3A/K4A and ΔStammer construct and performed initial DNA curtains experiments. C.K. expressed and purified His-TEV-ORF1p and performed fluorescence anisotropy and mass photometry for this construct. J.D.-L. expressed and purified ΔStammer-ORF1p. S.Z., F.E., M.F. and J.S. analyzed data. S.Z., F.E., L.J.H., and J.S. wrote the manuscript with input from all authors. S.Z., F.E., L.J.H. and J.S. designed the study. L.J.H and J.S. provided funding and supervised the project.

## Supplementary Figures

**Figure S1.**
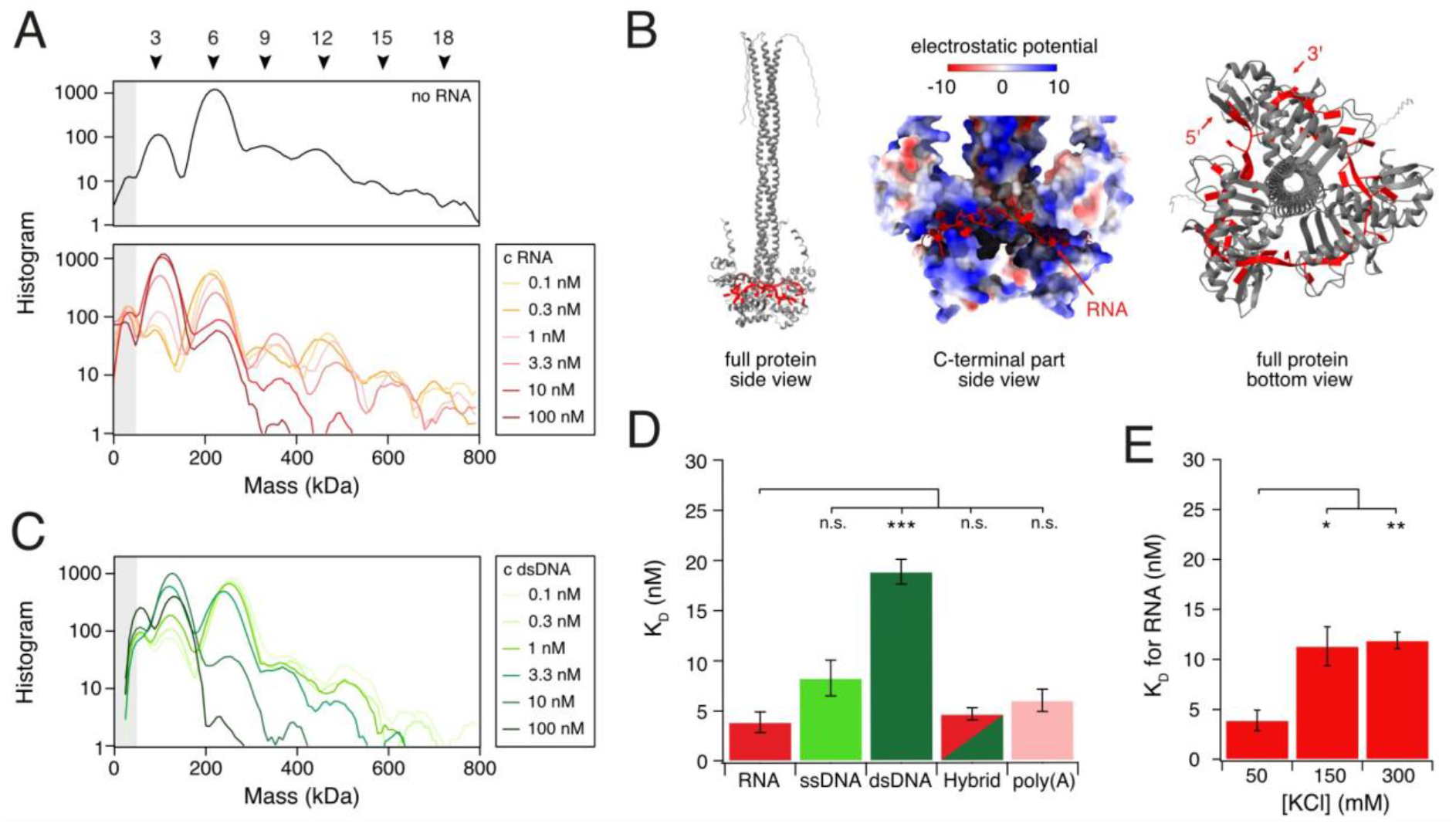
ORF1p binding to various nucleic acids. **(A)** Smoothed mass photometry histogram (log_10_ scale) of 300 nM ORF1p (top) and after binding to increasing concentrations of RNA oligonucleotide (bottom). Peaks for trimer and higher-order oligomers up to 18-mer could be assigned. At higher RNA concentrations, the trimer becomes dominant and fewer higher-order oligomers are present. **(B)** AlphaFold 3 model of ORF1p trimer (grey) binding to 30 nt RNA (red). Left: Full-length ORF1p. Middle: Zoom-in to the C-terminal domain showing the electrostatic surface potential of ORF1p. The RNA (red) is bound in the highly positively charged cleft (blue) between CTD and RRM. Right: Bottom-up view of the full-length protein. The RNA (red) is wrapped around the C-terminal part of the ORF1p trimer (grey). **(C)** Smoothed mass photometry histogram of 300 nM ORF1p binding to dsDNA. dsDNA binding induces hexamer dissociation similar to RNA. **(D)** Dissociation constants of ORF1p binding to different nucleic acid constructs determined by fluorescence anisotropy. n=3, error bars represent SEM. Significance was determined by Student’s t-test. **(E)** Dissociation constants for RNA binding of ORF1p at different salt concentrations.

**Figure S2.**
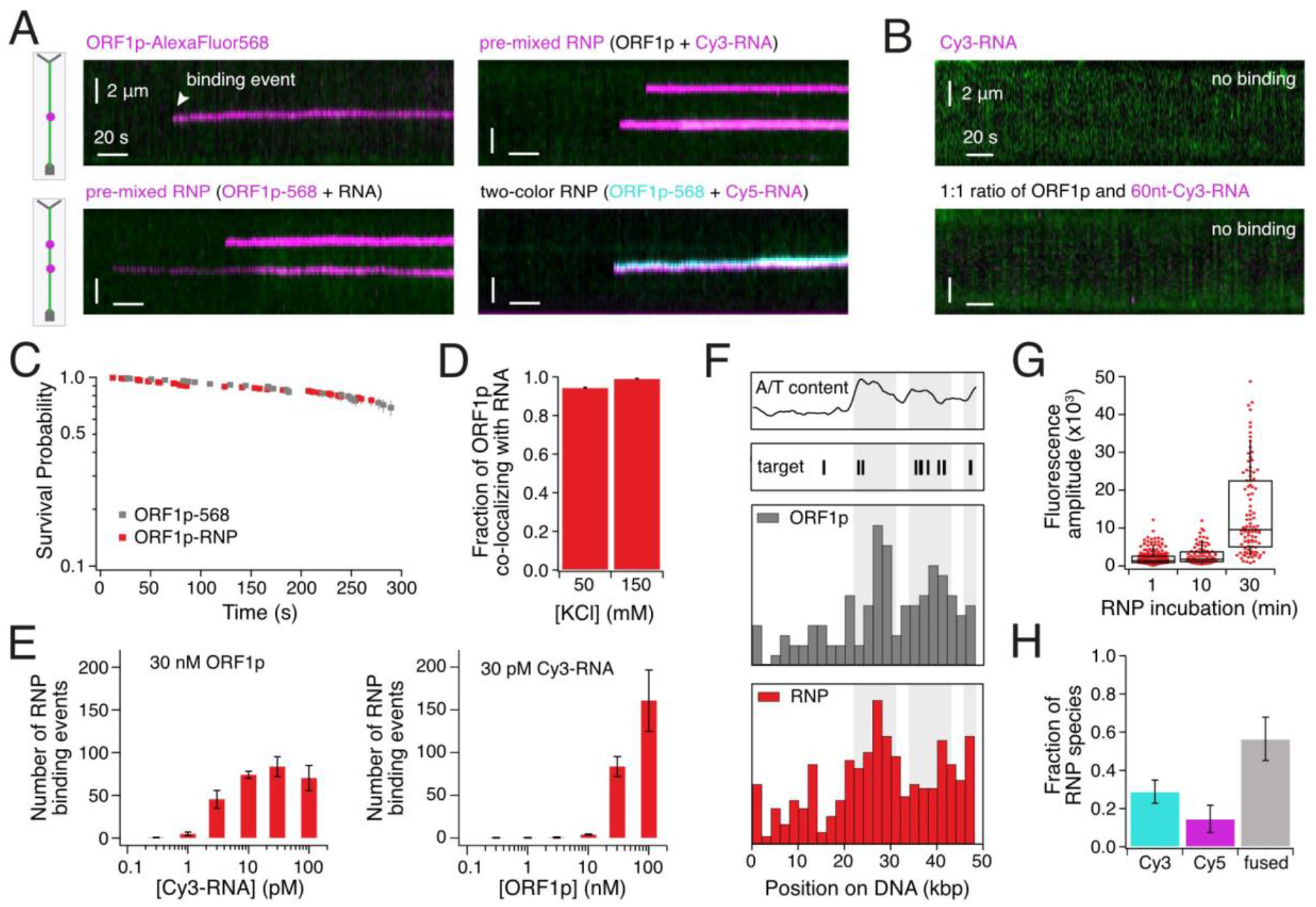
ORF1p-RNP binding characteristics on DNA curtains. **(A)** Representative kymograms of ORF1p or ORF1p-RNPs binding to DNA. **(B)** Neither long Cy3-labeled RNA alone (top, 30 pM of 2 kb RNA) nor 30 nM 60 nt Cy3-labeled RNA pre-mixed with 30 nM ORF1p (bottom) bind DNA. **(C)** Survival Plot of ORF1p or ORF1p-RNPs on DNA at 150 mM KCl. **(D)** In the presence of RNA, ORF1p binds to DNA as an RNP. RNPs in which both ORFp1 and RNA were labeled (ORF1p-AlexaFluor568 and Cy3-RNA), were counted during DNA binding in comparison to all binding events. The fraction of DNA-bound ORF1p that is also bound to Cy3-RNA is shown, n=3, error bars represent SEM. **(E)** Quantification of ORF1p-RNP binding events. Left: ORF1p concentration was kept constant at 30 nM and Cy3-RNA concentration was increased. Right: Cy3-RNA concentration was kept constant at 30 pM and ORF1p concentration was increased. n=3, error bars represent the SEM. **(F)** ORF1p prefers to bind to the A/T-rich DNA. The binding positions of ORF1p alone and ORF1p-RNPs on λ-DNA at 150 mM KCl are shown. The A/T-content of the DNA and the position of LINE-1 target sites (TTTTTAA) are plotted above the histograms. **(G)** With longer incubation times, RNPs become brighter. ORF1p-RNPs (30 nM ORF1p + 30 pM Cy3-RNA) were incubated for different lengths of time prior to loading onto DNA curtains and their fluorescence amplitude was measured once bound to DNA. n=3. **(H)** More than 50% of the RNPs contain at least two RNAs. RNPs were formed from ORF1p and RNA molecules labeled with one of two different dyes. Afterwards, the two samples were mixed and loaded on DNA curtains. The fraction of RNPs that contain both colors is shown, n=3, error bars represent the SEM.

**Figure S3.**
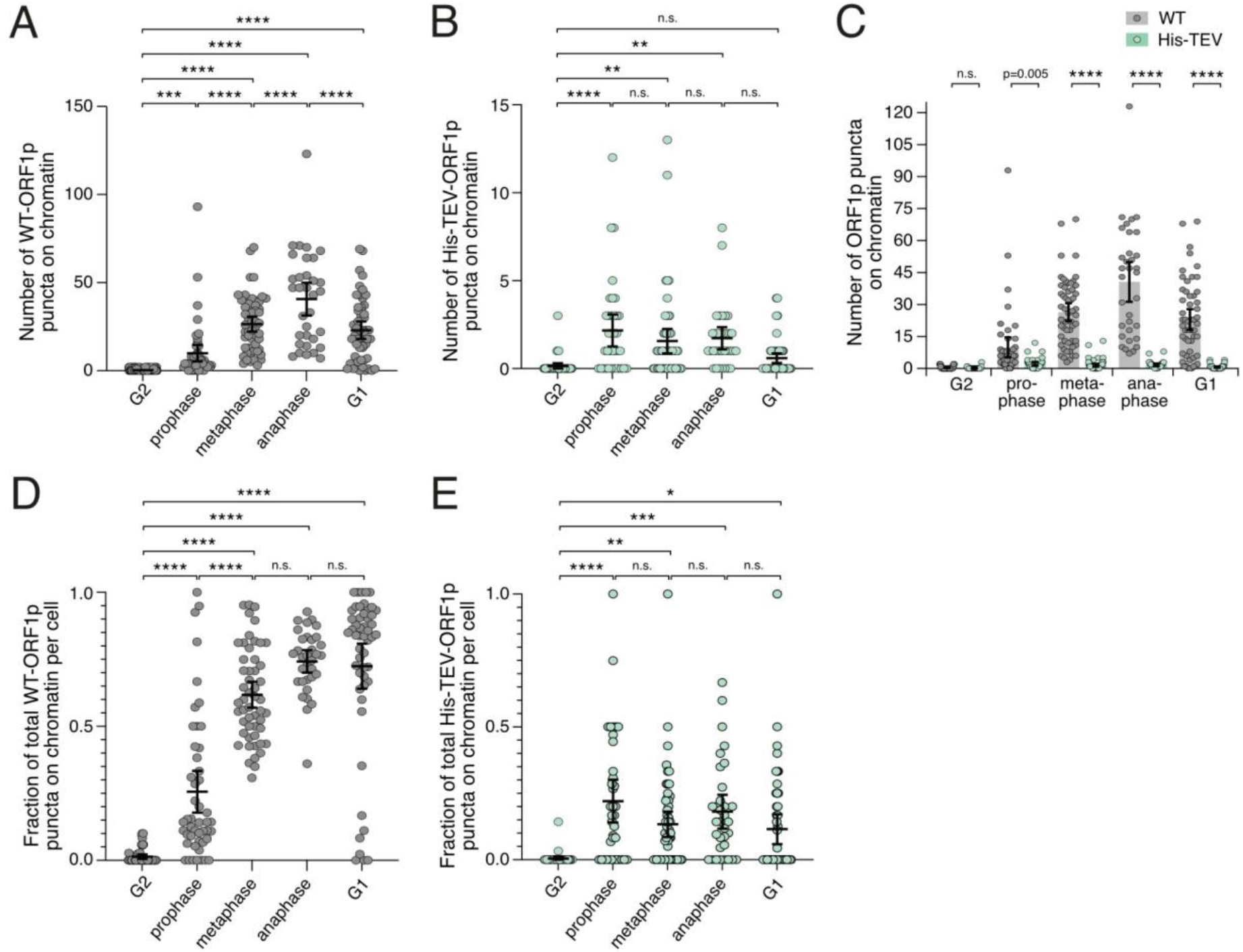
Puncta dynamics for WT-ORF1p and ORF1p mutants. **(A)** Absolute number of WT-ORF1p puncta colocalized with chromatin increases from prometaphase to anaphase. Significance determined by one-way ANOVA. N>30 per cell cycle stage, n=3. **(B)** Absolute number of His-TEV-ORF1p puncta colocalized with chromatin slightly increases upon nuclear envelope breakdown but remains stable thereafter. Significance determined by one-way ANOVA. N>30 per cell cycle stage, n=3. **(C)** Comparison of absolute numbers of WT- and His-TEV-ORF1p puncta co-localized with chromatin. WT has a significantly greater number of puncta co-localized with chromatin than His-TEV. Significance determined by multiple Student’s t-tests. N>30 per cell cycle stage, n=3. **(D)** Fraction of WT-ORF1p puncta colocalized with chromatin per cell across the cell cycle. Significance determined by one-way ANOVA. N>30 per cell cycle stage, n=3. **(E)** Fraction of His-TEV-ORF1p puncta colocalized with chromatin per cell across the cell cycle. Significance determined by one-way ANOVA. N>30 per cell cycle stage, n=3.

## References

1. Boeke, J.D., Garfinkel, D.J., Styles, C.A., and Fink, G.R. (1985). Ty elements transpose through an RNA intermediate. Cell 40, 491–500. 10.1016/0092-8674(85)90197-7.

2. Moran, J.V., Holmes, S.E., Naas, T.P., DeBerardinis, R.J., Boeke, J.D., and Kazazian, H.H. (1996). High frequency retrotransposition in cultured mammalian cells. Cell 87, 917–927. 10.1016/s0092-8674(00)81998-4.

3. Furano, A.V. (2000). The biological properties and evolutionary dynamics of mammalian LINE-1 retrotransposons. In Progress in Nucleic Acid Research and Molecular Biology (Academic Press), pp. 255–294. 10.1016/S0079-6603(00)64007-2.

4. Lander, E.S., Linton, L.M., Birren, B., Nusbaum, C., Zody, M.C., Baldwin, J., Devon, K., Dewar, K., Doyle, M., FitzHugh, W., et al. (2001). Initial sequencing and analysis of the human genome. Nature 409, 860–921. 10.1038/35057062.

5. Boissinot, S., Chevret, P., and Furano, A.V. (2000). L1 (LINE-1) Retrotransposon Evolution and Amplification in Recent Human History. Molecular Biology and Evolution 17, 915–928. 10.1093/oxfordjournals.molbev.a026372.

6. Boissinot, S., Entezam, A., Young, L., Munson, P.J., and Furano, A.V. (2004). The Insertional History of an Active Family of L1 Retrotransposons in Humans. Genome Res 14, 1221–1231. 10.1101/gr.2326704.

7. Boissinot, S., Davis, J., Entezam, A., Petrov, D., and Furano, A.V. (2006). Fitness cost of LINE-1 (L1) activity in humans. Proceedings of the National Academy of Sciences 103, 9590–9594. 10.1073/pnas.0603334103.

8. Chen, J.-M., Stenson, P.D., Cooper, D.N., and Férec, C. (2005). A systematic analysis of LINE-1 endonuclease-dependent retrotranspositional events causing human genetic disease. Hum Genet 117, 411–427. 10.1007/s00439-005-1321-0.

9. Beck, C.R., Garcia-Perez, J.L., Badge, R.M., and Moran, J.V. (2011). LINE-1 Elements in Structural Variation and Disease. Annu Rev Genomics Hum Genet 12, 187–215. 10.1146/annurev-genom-082509-141802.

10. Scott, A.F., Schmeckpeper, B.J., Abdelrazik, M., Comey, C.T., O’Hara, B., Rossiter, J.P., Cooley, T., Heath, P., Smith, K.D., and Margolet, L. (1987). Origin of the human L1 elements: Proposed progenitor genes deduced from a consensus DNA sequence. Genomics 1, 113–125. 10.1016/0888-7543(87)90003-6.

11. Mita, P., Wudzinska, A., Sun, X., Andrade, J., Nayak, S., Kahler, D.J., Badri, S., LaCava, J., Ueberheide, B., Yun, C.Y., et al. (2018). LINE-1 protein localization and functional dynamics during the cell cycle. eLife 7, e30058. 10.7554/eLife.30058.

12. Taylor, M.S., LaCava, J., Mita, P., Molloy, K.R., Huang, C.R.L., Li, D., Adney, E.M., Jiang, H., Burns, K.H., Chait, B.T., et al. (2013). Affinity Proteomics Reveals Human Host Factors Implicated in Discrete Stages of LINE-1 Retrotransposition. Cell 155, 1034–1048. 10.1016/j.cell.2013.10.021.

13. Feng, Q., Moran, J.V., Kazazian, H.H., and Boeke, J.D. (1996). Human L1 retrotransposon encodes a conserved endonuclease required for retrotransposition. Cell 87, 905–916. 10.1016/s0092-8674(00)81997-2.

14. Martin, S.L., Cruceanu, M., Branciforte, D., Wai-lun Li, P., Kwok, S.C., Hodges, R.S., and Williams, M.C. (2005). LINE-1 Retrotransposition Requires the Nucleic Acid Chaperone Activity of the ORF1 Protein. Journal of Molecular Biology 348, 549–561. 10.1016/j.jmb.2005.03.003.

15. Martin, S.L., Bushman, D., Wang, F., Li, P.W.-L., Walker, A., Cummiskey, J., Branciforte, D., and Williams, M.C. (2008). A single amino acid substitution in ORF1 dramatically decreases L1 retrotransposition and provides insight into nucleic acid chaperone activity. Nucleic Acids Res 36, 5845–5854. 10.1093/nar/gkn554.

16. Martin, S.L. (2010). Nucleic acid chaperone properties of ORF1p from the non-LTR retrotransposon, LINE-1. RNA Biol 7, 706–711. 10.4161/rna.7.6.13766.

17. Kolosha, V.O., and Martin, S.L. (1997). In vitro properties of the first ORF protein from mouse LINE-1 support its role in ribonucleoprotein particle formation during retrotransposition. Proc Natl Acad Sci U S A 94, 10155–10160.

18. Hohjoh, H., and Singer, M.F. (1996). Cytoplasmic ribonucleoprotein complexes containing human LINE-1 protein and RNA. The EMBO Journal 15, 630–639. 10.1002/j.1460-2075.1996.tb00395.x.

19. Sil, S., Keegan, S., Ettefa, F., Denes, L.T., Boeke, J.D., and Holt, L.J. (2023). Condensation of LINE-1 is critical for retrotransposition. eLife 12, e82991. 10.7554/eLife.82991.

20. Newton, J.C., Naik, M.T., Li, G.Y., Murphy, E.L., Fawzi, N.L., Sedivy, J.M., and Jogl, G. (2021). Phase separation of the LINE-1 ORF1 protein is mediated by the N-terminus and coiled-coil domain. Biophysical Journal 120, 2181–2191. 10.1016/j.bpj.2021.03.028.

21. Martin, S.L., Branciforte, D., Keller, D., and Bain, D.L. (2003). Trimeric structure for an essential protein in L1 retrotransposition. Proc Natl Acad Sci U S A 100, 13815–13820. 10.1073/pnas.2336221100.

22. Martin, S.L. (2006). The ORF1 Protein Encoded by LINE-1: Structure and Function During L1 Retrotransposition. J Biomed Biotechnol 2006, 45621. 10.1155/JBB/2006/45621.

23. Basame, S., Wai-lun Li, P., Howard, G., Branciforte, D., Keller, D., and Martin, S.L. (2006). Spatial Assembly and RNA Binding Stoichiometry of a LINE-1 Protein Essential for Retrotransposition. Journal of Molecular Biology 357, 351–357. 10.1016/j.jmb.2005.12.063.

24. Khazina, E., and Weichenrieder, O. (2009). Non-LTR retrotransposons encode noncanonical RRM domains in their first open reading frame. Proceedings of the National Academy of Sciences 106, 731–736. 10.1073/pnas.0809964106.

25. Erdel, F., and Rippe, K. (2018). Formation of Chromatin Subcompartments by Phase Separation. Biophysical Journal 114, 2262–2270. 10.1016/j.bpj.2018.03.011.

26. Wiedner, H.J., and Giudice, J. (2021). It’s not just a phase: function and characteristics of RNA-binding proteins in phase separation. Nat Struct Mol Biol 28, 465–473. 10.1038/s41594-021-00601-w.

27. Martin, E.W., and Holehouse, A.S. (2020). Intrinsically disordered protein regions and phase separation: sequence determinants of assembly or lack thereof. Emerging topics in life sciences 4. 10.1042/ETLS20190164.

28. Khazina, E., and Weichenrieder, O. (2018). Human LINE-1 retrotransposition requires a metastable coiled coil and a positively charged N-terminus in L1ORF1p. eLife 7, e34960. 10.7554/eLife.34960.

29. Goodier, J.L., Zhang, L., Vetter, M.R., and Kazazian, H.H. (2007). LINE-1 ORF1 Protein Localizes in Stress Granules with Other RNA-Binding Proteins, Including Components of RNA Interference RNA-Induced Silencing Complex. Mol Cell Biol 27, 6469–6483. 10.1128/MCB.00332-07.

30. Lupas, A. (1996). Prediction and analysis of coiled-coil structures. In Methods in Enzymology Computer Methods for Macromolecular Sequence Analysis. (Academic Press), pp. 513–525. 10.1016/S0076-6879(96)66032-7.

31. Januszyk, K., Li, P.W., Villareal, V., Branciforte, D., Wu, H., Xie, Y., Feigon, J., Loo, J.A., Martin, S.L., and Clubb, R.T. (2007). Identification and Solution Structure of a Highly Conserved C-terminal Domain within ORF1p Required for Retrotransposition of Long Interspersed Nuclear Element-1. Journal of Biological Chemistry 282, 24893–24904. 10.1074/jbc.M702023200.

32. Khazina, E., Truffault, V., Büttner, R., Schmidt, S., Coles, M., and Weichenrieder, O. (2011). Trimeric structure and flexibility of the L1ORF1 protein in human L1 retrotransposition. Nat Struct Mol Biol 18, 1006–1014. 10.1038/nsmb.2097.

33. Callahan, K.E., Hickman, A.B., Jones, C.E., Ghirlando, R., and Furano, A.V. (2012). Polymerization and nucleic acid-binding properties of human L1 ORF1 protein. Nucleic Acids Res 40, 813–827. 10.1093/nar/gkr728.

34. Naufer, M.N., Callahan, K.E., Cook, P.R., Perez-Gonzalez, C.E., Williams, M.C., and Furano, A.V. (2016). L1 retrotransposition requires rapid ORF1p oligomerization, a novel coiled coil-dependent property conserved despite extensive remodeling. Nucleic Acids Res 44, 281–293. 10.1093/nar/gkv1342.

35. Kolosha, V.O., and Martin, S.L. (2003). High-affinity, Non-sequence-specific RNA Binding by the Open Reading Frame 1 (ORF1) Protein from Long Interspersed Nuclear Element 1 (LINE-1). Journal of Biological Chemistry 278, 8112–8117. 10.1074/jbc.M210487200.

36. Cashen, B.A., Naufer, M.N., Morse, M., Jones, C.E., Williams, M.C., and Furano, A.V. (2022). The L1-ORF1p coiled coil enables formation of a tightly compacted nucleic acid-bound complex that is associated with retrotransposition. Nucleic Acids Res 50, 8690–8699. 10.1093/nar/gkac628.

37. Greene, E.C., Wind, S., Fazio, T., Gorman, J., and Visnapuu, M.-L. (2010). DNA Curtains for High-Throughput Single-Molecule Optical Imaging. Methods Enzymol 472, 293–315. 10.1016/S0076-6879(10)72006-1.

38. Young, G., Hundt, N., Cole, D., Fineberg, A., Andrecka, J., Tyler, A., Olerinyova, A., Ansari, A., Marklund, E.G., Collier, M.P., et al. (2018). Quantitative mass imaging of single molecules. Science 360, 423–427. 10.1126/science.aar5839.

39. Ortega-Arroyo, J., and Kukura, P. (2012). Interferometric scattering microscopy (iSCAT): new frontiers in ultrafast and ultrasensitive optical microscopy. Phys Chem Chem Phys 14, 15625–15636. 10.1039/c2cp41013c.

40. Cole, D., Young, G., Weigel, A., Sebesta, A., and Kukura, P. (2017). Label-Free Single-Molecule Imaging with Numerical-Aperture-Shaped Interferometric Scattering Microscopy. ACS Photonics 4, 211–216. 10.1021/acsphotonics.6b00912.

41. Abramson, J., Adler, J., Dunger, J., Evans, R., Green, T., Pritzel, A., Ronneberger, O., Willmore, L., Ballard, A.J., Bambrick, J., et al. (2024). Accurate structure prediction of biomolecular interactions with AlphaFold 3. Nature 630, 493–500. 10.1038/s41586-024-07487-w.

42. Rajagopalan, M., Balasubramanian, S., and Ramaswamy, A. (2017). Insights into the RNA binding mechanism of human L1-ORF1p: a molecular dynamics study. Mol. BioSyst. 13, 1728–1743. 10.1039/C7MB00358G.

43. Zernia, S., and Stigler, J. (2024). DNA curtains for studying phase separation mechanisms of DNA-organizing proteins. In Methods in Cell Biology (Academic Press), pp. 95–108. 10.1016/bs.mcb.2023.02.006.

44. Schorn, A.J., and Martienssen, R. (2019). Getting in LINE with Replication. Molecular Cell 74, 415–417. 10.1016/j.molcel.2019.04.023.

45. Jurka, J. (1997). Sequence patterns indicate an enzymatic involvement in integration of mammalian retroposons. Proceedings of the National Academy of Sciences 94, 1872–1877. 10.1073/pnas.94.5.1872.

46. Flasch, D.A., Macia, Á., Sánchez, L., Ljungman, M., Heras, S.R., García-Pérez, J.L., Wilson, T.E., and Moran, J.V. (2019). Genome-wide de novo L1 retrotransposition connects endonuclease activity with replication. Cell 177, 837. 10.1016/j.cell.2019.02.050.

47. Thawani, A., Ariza, A.J.F., Nogales, E., and Collins, K. (2024). Template and target-site recognition by human LINE-1 in retrotransposition. Nature 626, 186–193. 10.1038/s41586-023-06933-5.

48. Wu, D., and Piszczek, G. (2021). Standard Protocol for Mass Photometry Experiments. Eur Biophys J 50, 403–409. 10.1007/s00249-021-01513-9.

49. Castello, A., Fischer, B., Eichelbaum, K., Horos, R., Beckmann, B.M., Strein, C., Davey, N.E., Humphreys, D.T., Preiss, T., Steinmetz, L.M., et al. (2012). Insights into RNA Biology from an Atlas of Mammalian mRNA-Binding Proteins. Cell 149, 1393–1406. 10.1016/j.cell.2012.04.031.

50. Hentze, M.W., Castello, A., Schwarzl, T., and Preiss, T. (2018). A brave new world of RNA-binding proteins. Nat Rev Mol Cell Biol 19, 327–341. 10.1038/nrm.2017.130.

51. Protter, D.S.W., Rao, B.S., Van Treeck, B., Lin, Y., Mizoue, L., Rosen, M.K., and Parker, R. (2018). Intrinsically Disordered Regions Can Contribute Promiscuous Interactions to RNP Granule Assembly. Cell Rep 22, 1401–1412. 10.1016/j.celrep.2018.01.036.

52. Ottoz, D.S.M., and Berchowitz, L.E. (2020). The role of disorder in RNA binding affinity and specificity. Open Biol 10, 200328. 10.1098/rsob.200328.

53. Los, G.V., Encell, L.P., McDougall, M.G., Hartzell, D.D., Karassina, N., Zimprich, C., Wood, M.G., Learish, R., Ohana, R.F., Urh, M., et al. (2008). HaloTag: A Novel Protein Labeling Technology for Cell Imaging and Protein Analysis. ACS Chem. Biol. 3, 373–382. 10.1021/cb800025k.

54. Grimm, J.B., English, B.P., Chen, J., Slaughter, J.P., Zhang, Z., Revyakin, A., Patel, R., Macklin, J.J., Normanno, D., Singer, R.H., et al. (2015). A general method to improve fluorophores for live-cell and single-molecule microscopy. Nat Methods 12, 244–250. 10.1038/nmeth.3256.

55. Ostertag, E.M., Luning Prak, E.T., DeBerardinis, R.J., Moran, J.V., and Kazazian, H.H. (2000). Determination of L1 retrotransposition kinetics in cultured cells. Nucleic Acids Res 28, 1418–1423.

56. An, W., Dai, L., Niewiadomska, A.M., Yetil, A., O’Donnell, K.A., Han, J.S., and Boeke, J.D. (2011). Characterization of a synthetic human LINE-1 retrotransposon ORFeus-Hs. Mobile DNA 2, 2. 10.1186/1759-8753-2-2.

57. Wei, W., Gilbert, N., Ooi, S.L., Lawler, J.F., Ostertag, E.M., Kazazian, H.H., Boeke, J.D., and Moran, J.V. (2001). Human L1 Retrotransposition: cisPreference versus trans Complementation. Molecular and Cellular Biology 21, 1429–1439. 10.1128/MCB.21.4.1429-1439.2001.

58. Khong, A., and Parker, R. (2020). The landscape of eukaryotic mRNPs. RNA 26, 229–239. 10.1261/rna.073601.119.

59. Tauber, D., Tauber, G., and Parker, R. (2020). Mechanisms and Regulation of RNA Condensation in RNP Granule Formation. Trends Biochem Sci 45, 764–778. 10.1016/j.tibs.2020.05.002.

60. De Cecco, M., Ito, T., Petrashen, A.P., Elias, A.E., Skvir, N.J., Criscione, S.W., Caligiana, A., Brocculi, G., Adney, E.M., Boeke, J.D., et al. (2019). L1 drives IFN in senescent cells and promotes age-associated inflammation. Nature 566, 73–78. 10.1038/s41586-018-0784-9.

61. Hu, S., Li, J., Xu, F., Mei, S., Duff, Y.L., Yin, L., Pang, X., Cen, S., Jin, Q., Liang, C., et al. (2015). SAMHD1 Inhibits LINE-1 Retrotransposition by Promoting Stress Granule Formation. PLOS Genetics 11, e1005367. 10.1371/journal.pgen.1005367.

62. Arora, R., Bodak, M., Penouty, L., Hackman, C., and Ciaudo, C. (2022). Sequestration of LINE-1 in cytosolic aggregates by MOV10 restricts retrotransposition. EMBO reports 23, e54458. 10.15252/embr.202154458.

63. Kosinski, J., Mosalaganti, S., von Appen, A., Teimer, R., DiGuilio, A.L., Wan, W., Bui, K.H., Hagen, W.J.H., Briggs, J.A.G., Glavy, J.S., et al. (2016). Molecular architecture of the inner ring scaffold of the human nuclear pore complex. Science 352, 363–365. 10.1126/science.aaf0643.

64. Lin, D.H., Stuwe, T., Schilbach, S., Rundlet, E.J., Perriches, T., Mobbs, G., Fan, Y., Thierbach, K., Huber, F.M., Collins, L.N., et al. (2016). Architecture of the nuclear pore complex symmetric core. Science 352, aaf1015. 10.1126/science.aaf1015.

65. Suzuki, Y., and Craigie, R. (2007). The road to chromatin — nuclear entry of retroviruses. Nat Rev Microbiol 5, 187–196. 10.1038/nrmicro1579.

66. Booth, D.G., and Earnshaw, W.C. (2017). Ki-67 and the Chromosome Periphery Compartment in Mitosis. Trends in Cell Biology 27, 906–916. 10.1016/j.tcb.2017.08.001.

67. Cuylen-Haering, S., Petrovic, M., Hernandez-Armendariz, A., Schneider, M.W.G., Samwer, M., Blaukopf, C., Holt, L.J., and Gerlich, D.W. (2020). Chromosome clustering by Ki-67 excludes cytoplasm during nuclear assembly. Nature 587, 285–290. 10.1038/s41586-020-2672-3.

68. Booth, D.G., Takagi, M., Sanchez-Pulido, L., Petfalski, E., Vargiu, G., Samejima, K., Imamoto, N., Ponting, C.P., Tollervey, D., Earnshaw, W.C., et al. (2014). Ki-67 is a PP1-interacting protein that organises the mitotic chromosome periphery. eLife 3, e01641. 10.7554/eLife.01641.

69. Yamazaki, H., Takagi, M., Kosako, H., Hirano, T., and Yoshimura, S.H. (2022). Cell cycle-specific phase separation regulated by protein charge blockiness. Nat Cell Biol 24, 625–632. 10.1038/s41556-022-00903-1.

70. Molitor, T.P., and Traktman, P. (2014). Depletion of the protein kinase VRK1 disrupts nuclear envelope morphology and leads to BAF retention on mitotic chromosomes. Mol Biol Cell 25, 891–903. 10.1091/mbc.E13-10-0603.

71. Martin, S.L., and Bushman, F.D. (2001). Nucleic Acid Chaperone Activity of the ORF1 Protein from the Mouse LINE-1 Retrotransposon. Mol Cell Biol 21, 467–475. 10.1128/MCB.21.2.467-475.2001.

72. Evans, J.D., Peddigari, S., Chaurasiya, K.R., Williams, M.C., and Martin, S.L. (2011). Paired mutations abolish and restore the balanced annealing and melting activities of ORF1p that are required for LINE-1 retrotransposition. Nucleic Acids Res 39, 5611–5621. 10.1093/nar/gkr171.

73. Rief, M., Clausen-Schaumann, H., and Gaub, H. (1999). Sequence-dependent mechanics of single DNA molecules. Nature structural biology 6. 10.1038/7582.

